# Microglia deficiency accelerates prion disease but does not enhance prion accumulation in the brain

**DOI:** 10.1101/2021.01.05.425436

**Authors:** Barry M. Bradford, Lynne I. McGuire, David A. Hume, Clare Pridans, Neil A. Mabbott

## Abstract

Prion diseases are transmissible, neurodegenerative disorders associated with misfolding of the prion protein. Previous studies show that reduction of microglia accelerates CNS prion disease and increases the accumulation of prions in the brain, suggesting that microglia provide neuroprotection by phagocytosing and destroying prions. In *Csf1r*^ΔFIRE^ mice, the deletion of an enhancer within *Csf1r* specifically blocks microglia development, however, their brains develop normally and show none of the deficits reported in other microglia-deficient models. *Csf1r*^ΔFIRE^ mice were used as a refined model in which to study the impact of microglia-deficiency on CNS prion disease. Although *Csf1r*^ΔFIRE^ mice succumbed to CNS prion disease much earlier than wild-type mice, the accumulation of prions in their brains was reduced. Instead, astrocytes displayed earlier, non-polarized reactive activation with enhanced synaptic pruning and unfolded protein responses. Our data suggest that rather than simply phagocytosing and destroying prions, the microglia instead provide host-protection during CNS prion disease and restrict the harmful activities of reactive astrocytes.

**Main points:** CNS prion disease is accelerated in mice completely lacking microglia. The rate of prion accumulation in the brain was unaltered in absence of microglia. Microglia provide host-protection during CNS prion disease independent of prion clearance.

## Introduction

The parenchymal macrophages of the central nervous system (CNS) are known as microglia (Rio-Hortega, 1919) and their proliferation and survival is dependent upon signaling via the colony stimulating factor 1 receptor (CSF1R) (Hume et al., 2020). Microglia have been attributed essential functions in the development and homeostasis of the CNS including synaptogenesis, neurogenesis and maturation of neuronal circuits (Prinz et al., 2019). However, mice with a *Csf1r* hypomorphic mutation (*Csf1r*^ΔFIRE^) (Rojo et al., 2019), with conditional *Csf1r* deletion (using Iba1-cre) (Nakayama et al., 2018) and rats with a *Csf1r* null mutation (Pridans et al., 2018) each lack microglia entirely but have normal CNS development. These findings indicate that developmental roles of microglia are redundant as studies reveal their functions can be carried out by other cells when microglia are absent (Guo et al., 2019, Damisah et al., 2020, Patkar et al., 2021). There is much greater evidence that microglia contribute to neuropathology (Prinz et al., 2019). Neurodegenerative diseases associated with mutations in microglia-expressed genes such as *CSF1R* in humans have been referred to as microgliopathies (Hume et al., 2020).

Prion diseases, or transmissible spongiform encephalopathies, are fatal progressive neurodegenerative diseases to which there are no cures. Infectious prions are considered to result from the misfolding of the host’s cellular prion protein (PrP^C^) into an abnormal disease-associated isoform (PrP^Sc^) (Prusiner, 1982). The accumulation of PrP^Sc^ within the brain is accompanied by the impairment of neuronal dendritic spines and synapse structures, glial cell activation, vacuolar (spongiform) degeneration and ultimately neurodegeneration. Inhibiting the proliferation and pro-inflammatory responses of microglia via CSF1R inhibition decelerated CNS prion disease (Gómez-Nicola et al., 2013). Conversely, the partial depletion or deficiency in microglia was reported to enhance the accumulation of prions in the brain and accelerate the onset of clinical disease (Zhu et al., 2016, Carroll et al., 2018). However, none of these studies resulted in 100% microglial ablation nor addressed the potential confounding effects of ablative cell death or bystander effects, such as impact upon other non-microglial CSF1R-sensitive mononuclear phagocyte populations. For example although the CSF1R-targeting kinase inhibitor PLX5622 has been widely used to ablate the microglia in the brain, such kinase inhibitors also impact peripheral CSF1R-dependent macrophages (Hume and MacDonald, 2012). Since the ablation of peripheral macrophages enhances prion accumulation in the secondary lymphoid tissues (Beringue et al., 2000, Maignien et al., 2005), effects on peripheral macrophage populations in the above studies also cannot be excluded.

To address the above concerns we investigated CNS prion disease in *Csf1r*^ΔFIRE^ mice which have a complete and specific lack of microglia in the brain but retain brain-associated macrophages (Rojo et al., 2019). We show that microglial-deficiency in *Csf1r*^ΔFIRE^ mice was associated with accelerated prion disease in the absence of increased PrP^Sc^ accumulation or prion-seeding activity. Instead, earlier astrocyte activation was associated with increased synaptic engulfment and unfolded protein responses without induction of genes associated with neurotoxic (A1) or neuroprotective (A2) reactive astrocyte polarization (Liddelow et al., 2017). These data indicate that microglia provide neuroprotection during CNS prion disease independently of PrP^Sc^ clearance, and restrict the harmful effects of reactive astrocyte activation. Identification of the mechanisms by which the microglia provide neuroprotection during CNS prion disease may reveal novel targets for therapeutic intervention in these and other neurodegenerative disorders.

## Materials and methods

### Ethics statement

Ethical approvals for the *in vivo* mouse experiments were obtained from The Roslin Institute’s and University of Edinburgh’s ethics committees. These experiments were also performed under the authority of a UK Home Office Project Licence and in accordance with the guidelines and regulations of the UK Home Office ‘Animals (scientific procedures) Act 1986’. Appropriate care was provided to minimise harm and suffering, and anaesthesia was administered where necessary. Mice were humanely culled at the end of the experiments by cervical dislocation.

### Mice

*Csf1r*^ΔFIRE/WT^ mice produced in-house (Rojo et al., 2019) were crossed to produce homozygous *Csf1r*^ΔFIRE^ (*Csf1r*^ΔFIRE/ΔFIRE^) or *Csf1r*^WT^ (*Csf1r*^WT/WT^) littermates. Offspring were genotyped as described (Rojo et al., 2019). Pups were weaned and co-housed under specific pathogen-free conditions. Food and water were provided ad libitum.

### Prion infection

Mice were infected at 10 weeks old via intracerebral injection with 20 µl of a 0.01% (weight/volume) brain homogenate prepared from mice terminally infected with ME7 scrapie prions. Mice were culled at the intervals indicated after exposure, or observed for signs of clinical prion disease as described elsewhere (Brown and Mabbott, 2014) and culled at a standard clinical end-point. Survival times were calculated as the interval between injection and positive clinical assessment of terminal prion disease.

### Gait analysis

Gait analysis was performed weekly using the CatWalkXT (Noldus) from 8 weeks of age until positive clinical assessment of prion disease. Uninfected mice of both genotype were monitored weekly from 8 to 30 weeks of age as controls.

### Neuropathological analysis

Clinical prion disease diagnosis was confirmed by histopathological assessment of vacuolation (spongiform pathology) in the brain. Coronal sections of paraffin-embedded brain tissue were cut at 6 µm thickness, de-paraffinized and stained with hematoxylin & eosin and scored for spongiform vacuolar degeneration as described previously (Fraser and Dickinson, 1967). For the construction of lesion profiles, sections were scored for the presence and severity (scale 0–5) of prion-disease-specific vacuolation in nine grey matter and three white matter areas: G1, dorsal medulla; G2, cerebellar cortex; G3, superior colliculus; G4, hypothalamus; G5, thalamus; G6, hippocampus; G7, septum; G8, retrosplenial and adjacent motor cortex; G9, cingulate and adjacent motor cortex; W1, inferior and middle cerebellar peduncles; W2, decussation of superior cerebellar peduncles; and W3, cerebellar peduncles.

### Immunohistochemistry

Paraffin-embedded sections (thickness 6 μm) were deparaffinized, pre-treated by autoclaving in distilled water at 121°C for 15 min, and for PrP-immunostaining immersed in 98% formic acid for 10 min, endogenous peroxidases were quenched by immersion in 4% H_2_0_2_ in methanol for 5 min. Sections were incubated overnight with primary antibodies (see Table 1). Primary antibody binding was detected using biotinylated goat anti-species specific antibodies (Jackson Immunoresearch, West Grove, PA) where necessary and visualized using the Elite ABC/HRP kit (Vector Laboratories, Peterborough, UK) and diaminobenzidine (DAB) between stringent washing steps. Sections were lightly counterstained with hematoxylin and imaged on a Nikon Ni.1 Brightfield Compound upright microscope, 4x/10x/20x/ air lenses, Zeiss 105c colour camera & Zen 2 software for image capture. For fluorescence immunohistochemistry primary antibodies were detected with species-specific Alexa-Fluor 488 or 594 conjugated secondary antibodies. Perk-P staining was detected using biotinylated goat anti-rabbit specific antibodies (Jackson Immunoresearch, West Grove, PA) and visualized using the Elite ABC/HRP kit (Vector Laboratories, Peterborough, UK) and Tyramide Alexa-Fluor488 (Biotium) and imaged on a Zeiss LSM 710 Confocal Microscope with 6 Laser Lines (405/458/488/514/543/633nm)/ 2 PMT’s + 32 channel Quasar detector. 10×/20×/40×1.3na oil/60×1.4na oil lenses using Zen Software.

**Table 1:**
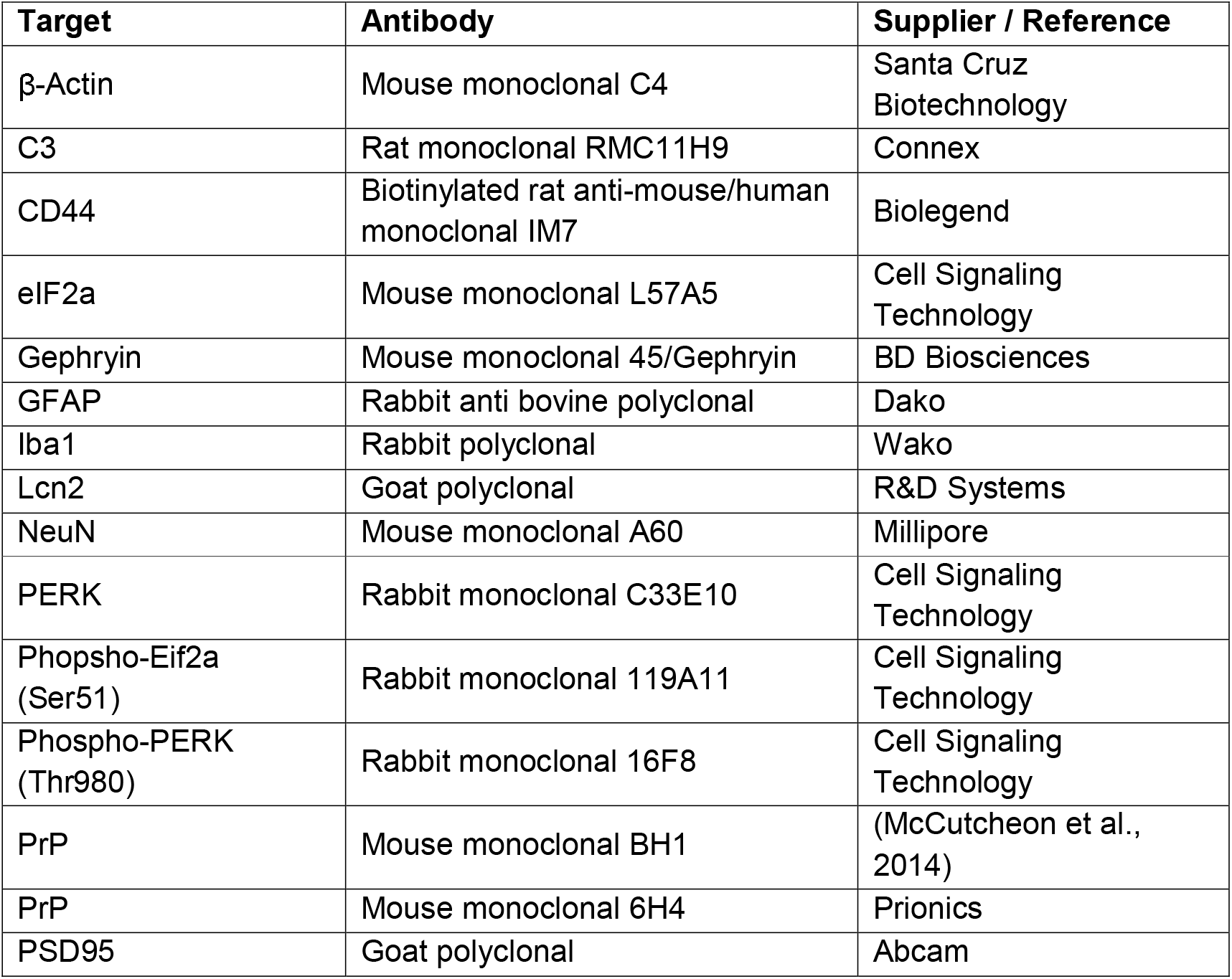
Primary antibodies.

### Western blot analysis

Brain homogenates (10% weight/volume) were prepared in NP40 lysis buffer (1% NP40, 0.5% sodium deoxycholate, 150 mM NaCl, 50 mM TrisHCl [pH 7.5]). For the detection of PrP^Sc^ a sample of homogenate was incubated at 37°C for 1 h with 20 µg/ml proteinase K (PK) and digestion halted by addition of 1 mM phenylmethylsulfonyl fluoride. Samples were denatured at 98°C for 15 min in 1x SDS sample buffer (Life Technologies) and separated via electrophoresis through 12% Tris-glycine polyacrylamide gels (Nupage, Life Technologies) and transferred to polyvinylidene difluoride PVDF membranes by semi-dry electroblotting. Primary antibodies (**Table 1**) were detected by horseradish peroxidase-conjugated goat anti-species specific antibody (Jackson Immunoresearch) and visualized via chemiluminescence (BM Chemiluminescent substrate kit, Roche, Burgess Hill, UK) as described previously (Bradford et al., 2017).

### Image analyses

Image analysis was performed using ImageJ software (http://imagej/nih.gov/ij) (Schneider et al., 2012). The magnitude of PrP^d^, GFAP and CD44 immunostaining on DAB stained sections was compared as previously described (Bradford et al., 2019). Briefly, the optical density (OD) values for immunostaining were calculated using ImageJ software following H-DAB deconvolution. Mean grey OD values were measured from DAB grayscale images (scaled 0–255) and expressed as a % relative intensity by dividing by the maximum value (255). Immunofluorescent images were analysed using ImageJ as previously described (McCulloch et al., 2011). Briefly intensity thresholds were applied and then the number of pixels of each colour counted and presented as a proportion of the total pixel area under analysis (% area coverage). The preferential co-localisation of fluorochromes were determined by comparison of the observed distribution of colors with those predicted by the null hypothesis that each element of positive staining was randomly and independently distributed. Values found to be significantly greater than the null hypothesis confirm significant co-localisation of fluorochromes. The assessment of relative synaptic pruning was calculated as the % of PSD95 staining co-localised with GFAP relative to total PSD95. Western blot images were subjected to densitometric analysed by ImageJ. Briefly lanes and bands were identified, threshold levels set and area under the curve measurements taken (pixels). For PrP^C^ and PrP^Sc^ relative expression levels were calculated as a percentage relative to a control normal brain PrP^C^ measurement.

### Real-Time quaking induced conversion (RT-QuiC)

Brain homogenates were diluted at 10−3 volume/volume in PBS. RT-QuIC reaction mix prepared as follows: 10 mM phosphate buffer (pH 7.4), 170 mM NaCl (total 300 mM including phosphate buffer), 0.1 mg/mL recombinant PrPc (Bank Vole 23–230, (Orrú et al., 2015) construct kindly provided by Byron Caughey), 10 μM Thioflavin-T (ThT), and 10 μM ethylenediaminetetraacetic acid tetrasodium salt (EDTA). Reactions were performed in quadruplicate. Aliquots of the reaction mix (98 μL) were loaded into each well of a black 96-well plate with a clear bottom (Thermo Scientific) and seeded with 2 μL of diluted brain homogenate. Samples were incubated in a FLUOstar® OMEGA plate reader (BMG LABTECH Ltd.) at 42°C for 80 h with intermittent shaking cycles: 1 min shake (double orbital, 700rpm), 1 min rest. Fluorescence measurements (450 nm excitation and 480 nm emission; bottom read), referred to as relative fluorescent units (rfu) were taken every 15 min. A baseline rfu of ∼38,000 for unseeded and initial BH seeded reactions were recorded, with saturation occurring at 260,000 rfu. All 4 quadruplicates of the 8 test samples, displayed a significant rise in rfu over time; a sample was considered “positive for PrP seeding” if replicates crossed a threshold of fluorescence set at 50,000 rfu based on the mean ± 10SD (36941 ± 8348) of the unseeded negative control samples analysed. The mean time for each quadruplicate reading to reach the 50,000 rfu threshold was calculated and plotted.

### Gene expression analysis via quantitative RT-PCR

Total RNA was isolated from brain using RNABee (AMSBio, Abingdon, UK) and RNeasy Mini kit (Qiagen). RNA was Dnase treated (Promega) to remove genomic DNA. Reverse transcription of polyA mRNA from 5 µg total DNA-free RNA was performed using Superscript First Strand Synthesis (Invitrogen) with Oligo-dT primers. Quantitative PCR (qPCR) were performed using SYBR master mix (Rox) (Roche) on an MX3005pro (Stratagene) using the primer sequences detailed (**Table 2**). Gene expression relative to naïve *Csf1r*^WT^ mice was calculated using the ΔΔCT method (Livak and Schmittgen, 2001) using *Rpl19* as a reference gene.

**Table 2:**
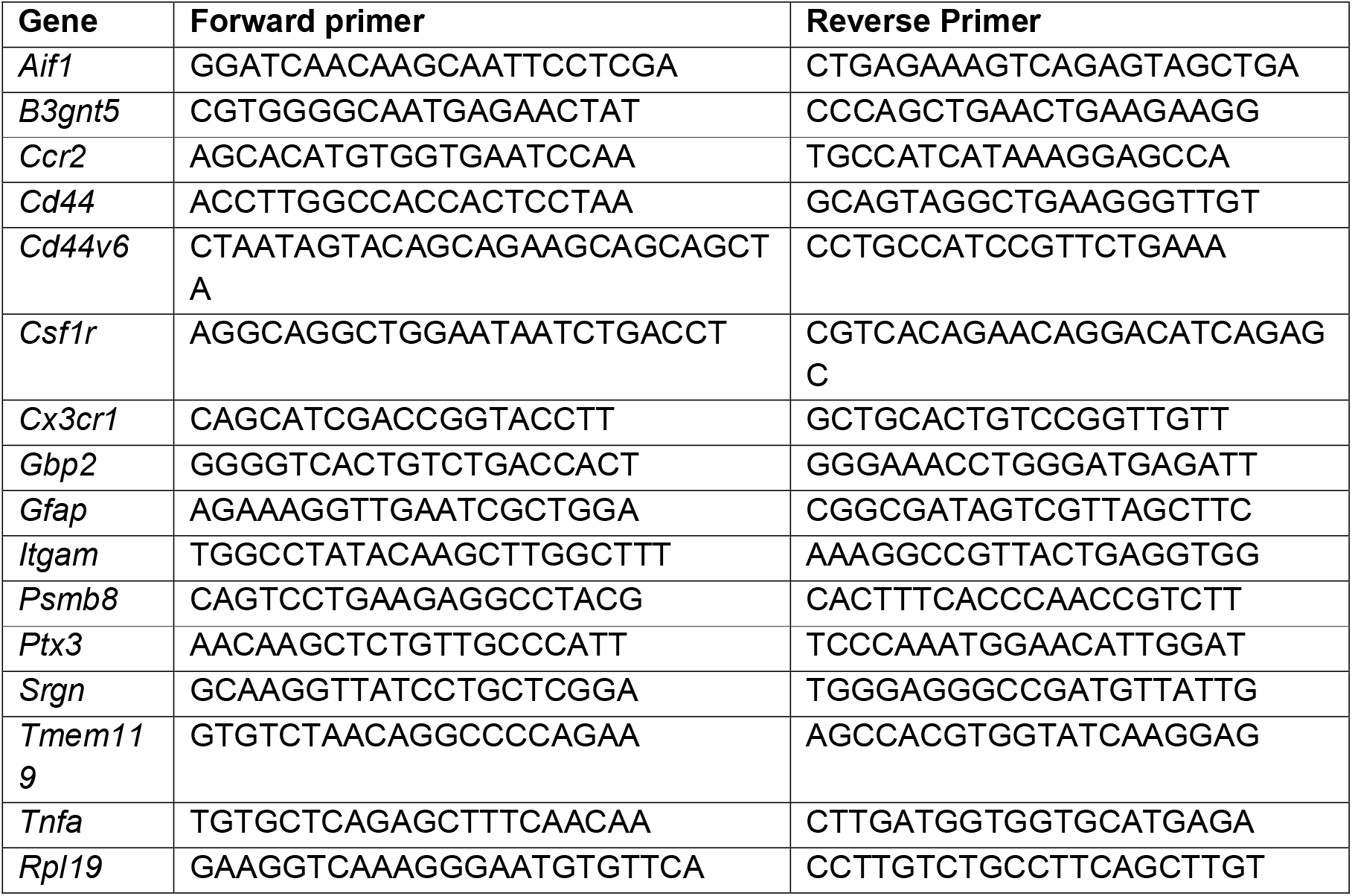
Oligonucleotide primers

### Data availability

The data that support the findings of this study are available from the corresponding author upon reasonable request.

### Statistical analyses

Statistical analysis was performed in GraphPad Prism 6.01 (GraphPad Software Inc.). Survival curve analysis performed by Log-rank [Mantel Cox] Test. Image and gene expression analyses performed by Student’s *t*-test (2 groups) or ANOVA (4 groups) results expressed as dot plots of individual animal observations with median values indicated (bar). CatWalkXT analysis performed using two-way ANOVA and expressed as group mean with 95% confidence interval. Values of *P*□<□0.05 were accepted as significant.

## Results

### *Csf1r*^ΔFIRE^ mice rapidly succumb to prion disease in the absence of microglia

To determine the role of microglia in prion disease, groups of homozygous microglia-deficient *Csf1r*^ΔFIRE^ transgenic mice and wild-type (*Csf1r*^WT^) littermate controls were injected intracerebrally (IC) with the ME7 strain of mouse adapted scrapie prions. As expected, all the *Csf1r*^WT^ mice displayed clinical signs of prion disease from approximately 140 days after injection and succumbed to terminal disease with a mean survival time of 167 ± 5 days. In *Csf1r*^ΔFIRE^ mice, clinical manifestations of prion disease were evident by 98 days after infection and progressed rapidly resulting in a mean survival time of 124 ± 2 days (**Figure 1A**).

**Figure 1.**
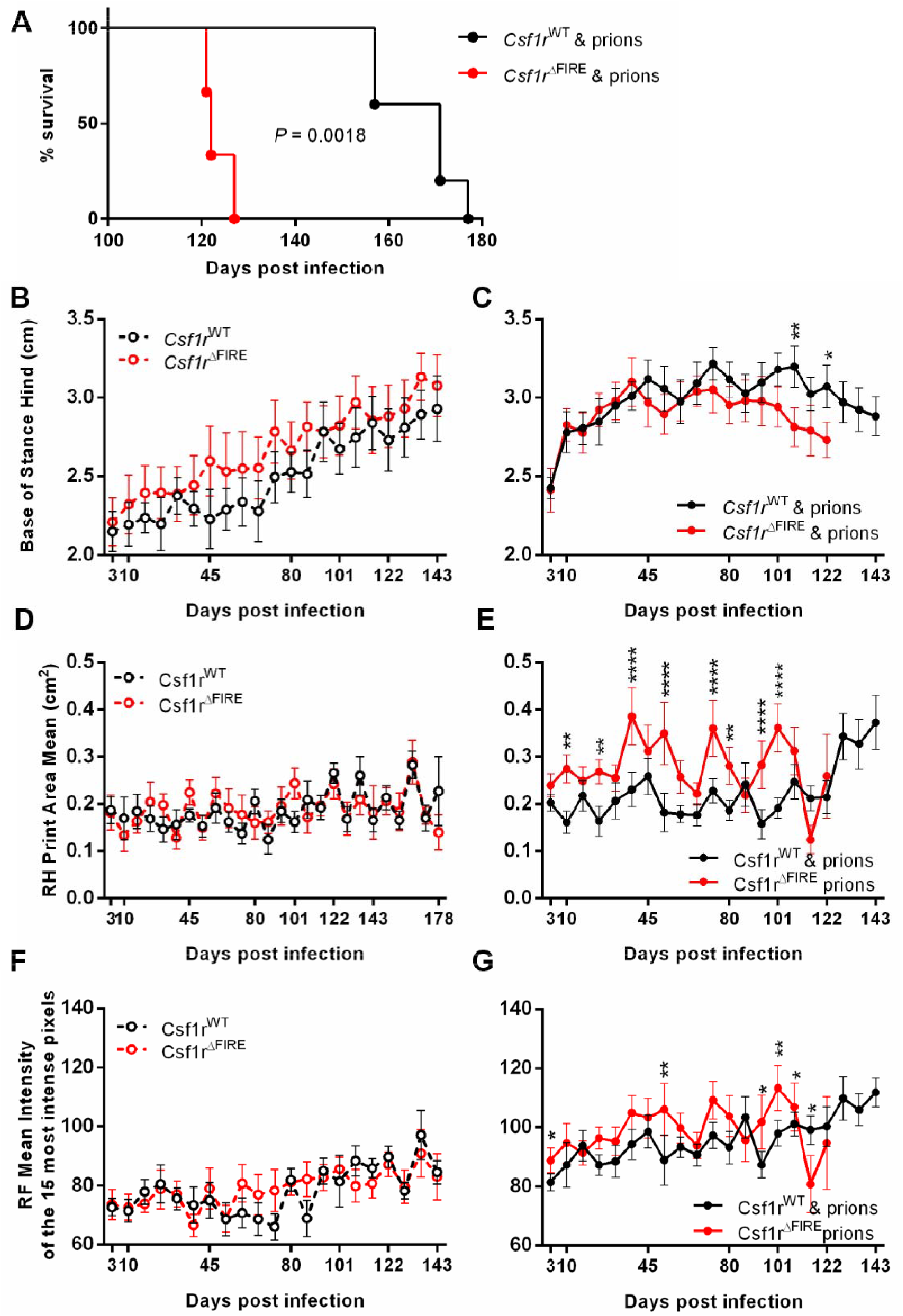
*Csf1r*^ΔFIRE^ mice rapidly succumb to prion disease. (A) Survival curve following intracerebral injection of ME7 prions into *Csf1r*^WT^ or *Csf1r*^ΔFIRE^ mice N=5-6. Log-rank Mantel Cox Test, *P* = 0.0018. (B) Catwalk XT automated gait analysis weekly assessment of hind base of stance in uninfected *Csf1r*^WT^ or *Csf1r*^ΔFIRE^ mice N=6-10. Points represent group mean and error bars 95% confidence interval. Two-way ANOVA. (C) Weekly assessment of hind base of stance in prion-infected *Csf1r*^WT^ or *Csf1r*^ΔFIRE^ mice N=6-10. Two-way ANOVA, Sidak’s multiple comparisons test \**P* < 0.05, \*\**P* < 0.005. (D) Weekly assessment of right hind (RH) paw print area in uninfected mice N=6-10. Two-way ANOVA. (E) Weekly assessment of right hind (RH) paw print area in prion-infected mice N=6-10. Two-way ANOVA, Sidak’s multiple comparisons test \*\**P* < 0.005, \*\*\*\**P* < 0.0001. (F) Weekly assessment of right front (RF) paw intensity in uninfected mice N=6-10. Two-way ANOVA. (G) Weekly assessment of right front (RF) paw intensity in prion-infected mice N=6-10. Two-way ANOVA, Sidak’s multiple comparisons test \**P* < 0.05, \*\**P* < 0.005.

### Longitudinal gait analysis during prion infection

CNS prion disease in mice is associated with profound motor-coordination disturbances (Heitzman and Corp, 1968). Since microglia are proposed to be involved in the development of the cerebellum and motor function (Kana et al., 2019), we used longitudinal gait analysis to determine whether microglia-deficiency affected the onset of motor disturbances during CNS prion disease (**Figure 1B-G**). Contrary to the published study, our analyses revealed no significant impact of the complete absence of microglia in the cerebellum of *Csf1r*^ΔFIRE^ mice on motor function analyzed at any time point in the absence of the prion challenge (**Figure 1B, D&F**).

As expected, various motor functions were rapidly impacted in prion disease. The base of stance (BOS, or distance between the hind paws) increased gradually with age in uninfected mice regardless of genotype (**Figure 1B**) but diverged by 10 days post infection (dpi) with prions and was maintained until 63 dpi (9 weeks) in *Csf1r*^ΔFIRE^ mice and 108 dpi (15 weeks) in *Csf1r*^WT^ mice (**Figure 1C**). At the onset of clinical signs of prion disease at 101 dpi) *Csf1r*^ΔFIRE^ mice were hyperactive and continued to perform Catwalk Gait analysis with ease until the terminal stage. In contrast, *Csf1r*^WT^ mice at the onset of clinical symptoms at 143 dpi were severely ataxic and unable to cross the Catwalk within the time-period required for data acquisition (**Figure 1C**).

Due to the potential effects of IC injection of prions into the right hemisphere on the contralateral paws, we analyzed the effects on footfall using only the unilateral right front and hind paws. In uninfected mice hind paw area remained unchanged with no statistically significant differences between *Csf1r*^WT^ and *Csf1r*^ΔFIRE^ mice (**Figure 1D)**. A significant increase in hind paw area was observed at 10 dpi in prion-infected *Csf1r*^ΔFIRE^ mice with a further large increase at 38 dpi. In contrast *Csf1r*^WT^ mice did not experience a large increase in right hind paw area until 129 dpi, 3 weeks before commencement of clinical signs, these data are indicative of a more rapid response to prion infection in *Csf1r*^ΔFIRE^ mice (**Figure 1E**).

In uninfected mice front paw intensity remained unchanged with no statistically significant differences between *Csf1r*^WT^ and *Csf1r*^ΔFIRE^ mice (**Figure 1F)**. Concurrent with changes in footprint area, footfall intensity was increased in the prion-infected mice (**Figure 1G**). Front footfall intensity increased significantly from 3 dpi in *Csf1r*^WT^ mice and this increase was maintained almost throughout the duration of the prion infection until the terminal stage. In contrast, footfall intensity in the prion-infected *Csf1r*^ΔFIRE^ mice commenced at 10 dpi and was maintained until onset of clinical symptoms at 101 dpi (**Figure 1G**).

### Detection of microgliosis in the prion-infected WT mice

The brains of terminal prion-infected *Csf1r*^WT^ mice displayed abundant, activated microglia (allograft inhibitory factor-1-positive [AIF1^+^] cells), whereas these cells and other potential AIF1^+^ CNS-infiltrating mononuclear phagocyte populations remained absent in uninfected and terminally-affected *Csf1r*^ΔFIRE^ mice (**Figure 2A&B**)(Rojo et al., 2019). RT-qPCR analysis confirmed that *Aif1* (**Figure 2C**) and *Csf1r* (**Figure 2D**) mRNA expression was significantly increased in the brains of terminally-affected *Csf1r*^WT^ mice when compared to uninfected controls, but remained almost undetectable in the brains of *Csf1r*^ΔFIRE^ mice even at the terminal stage of prion disease. Expression of other important microglia genes including *Itgam* (encoding CD11b; **Figure 2E**), *Cx3cr1* (**Figure 2F**) and *Tmem119* (**Figure 2G**) were also significantly increased in *Csf1r*^WT^ mice, but absent in *Csf1r*^ΔFIRE^ mice at the terminal stage of prion infection. The monocyte chemokine receptor *Ccr2* (**Figure 2H**) was significantly increased following prion infection in *Csf1r*^WT^ mice, but not in *Csf1r*^ΔFIRE^ mice. Together, these data show that onset of CNS prion disease was accelerated in *Csf1r*^ΔFIRE^ mice in the complete absence of microglia. Notably, the *Csf1r*^ΔFIRE^ mice are not monocyte-deficient but their monocytes lack CSF1R expression (Rojo et al, 2019). The IHC and expression profiling indicates that the *Csf1r*^ΔFIRE^ mutation also prevents monocyte recruitment into the injured brain.

**Figure 2.**
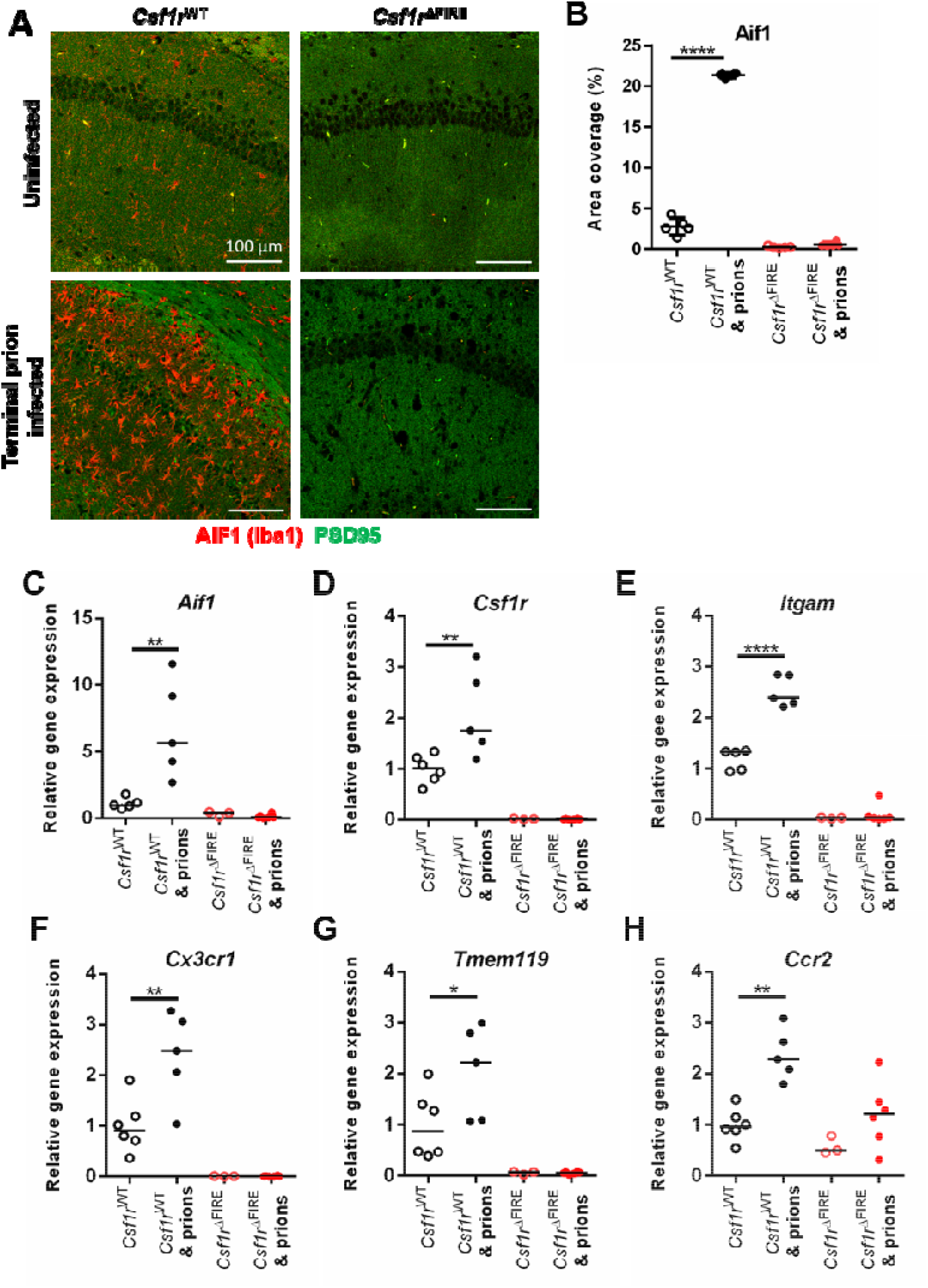
*Csf1r*^ΔFIRE^ mice succumb to prion disease in the absence of microglia. (A) Immunohistochemical assessment of AIF1 (red) and PSD95 (green) in hippocampus CA1 of uninfected or terminal prion infected *Csf1r*^WT^ or *Csf1r*^ΔFIRE^ mice. Scale bars = 100 µm. (B) AIF1 immunostaining quantitation expressed as % area coverage in hippocampus CA1. Points show individual mice as indicated, bar = median. ANOVA \*\*\*\**P* < 0.0001. (C) *Aif1* expression analysis via RT-qPCR on uninfected or terminal prion-infected brains from *Csf1r*^WT^ or *Csf1r*^ΔFIRE^ mice. Points show individual mice as indicated, bar = median. ANOVA \*\**P* < 0.005. (D) *Csf1r* expression analysis. ANOVA \*\**P* < 0.005. *(E) Itgam* expression analysis. ANOVA \*\*\*\**P* < 0.0001. (*F) Cx3cr1* expression analysis. ANOVA \*\**P* < 0.005. *(G) Tmem119* expression analysis. ANOVA \**P* < 0.05. (H) *Ccr2* expression analysis. ANOVA. \*\**P* < 0.01.

### Unaltered neuronal loss but reduced prion accumulation in the brains of microglia-deficient mice

Assessment of hippocampal CA1 pyramidal cells in hematoxylin and eosin stained brain sections (**Figure 3A**) revealed no difference in neuronal density (**Figure 3B**) or the frequency of pyknotic (apoptotic) neuronal nuclei between terminal prion-infected *Csf1r*^WT^ and *Csf1r*^ΔFIRE^ mice despite the difference in time of onset of pathology (**Figure 3C**). The prion-specific vacuolation was also comparable in most brain areas of *Csf1r*^WT^ and *Csf1r*^ΔFIRE^ terminal prion-infected mice, except for a significant reduction of vacuolation in the cerebellar cortex (G2), inferior and middle cerebellar peduncles (W1) and decussation of superior cerebellar peduncles (W2) of brains from prion-infected *Csf1r*^ΔFIRE^ mice (**Figure 3D**). This suggested the pathological impact of prion-infection upon the cerebellum was reduced in the *Csf1r*^ΔFIRE^ mice at the terminal stage of prion disease.

**Figure 3.**
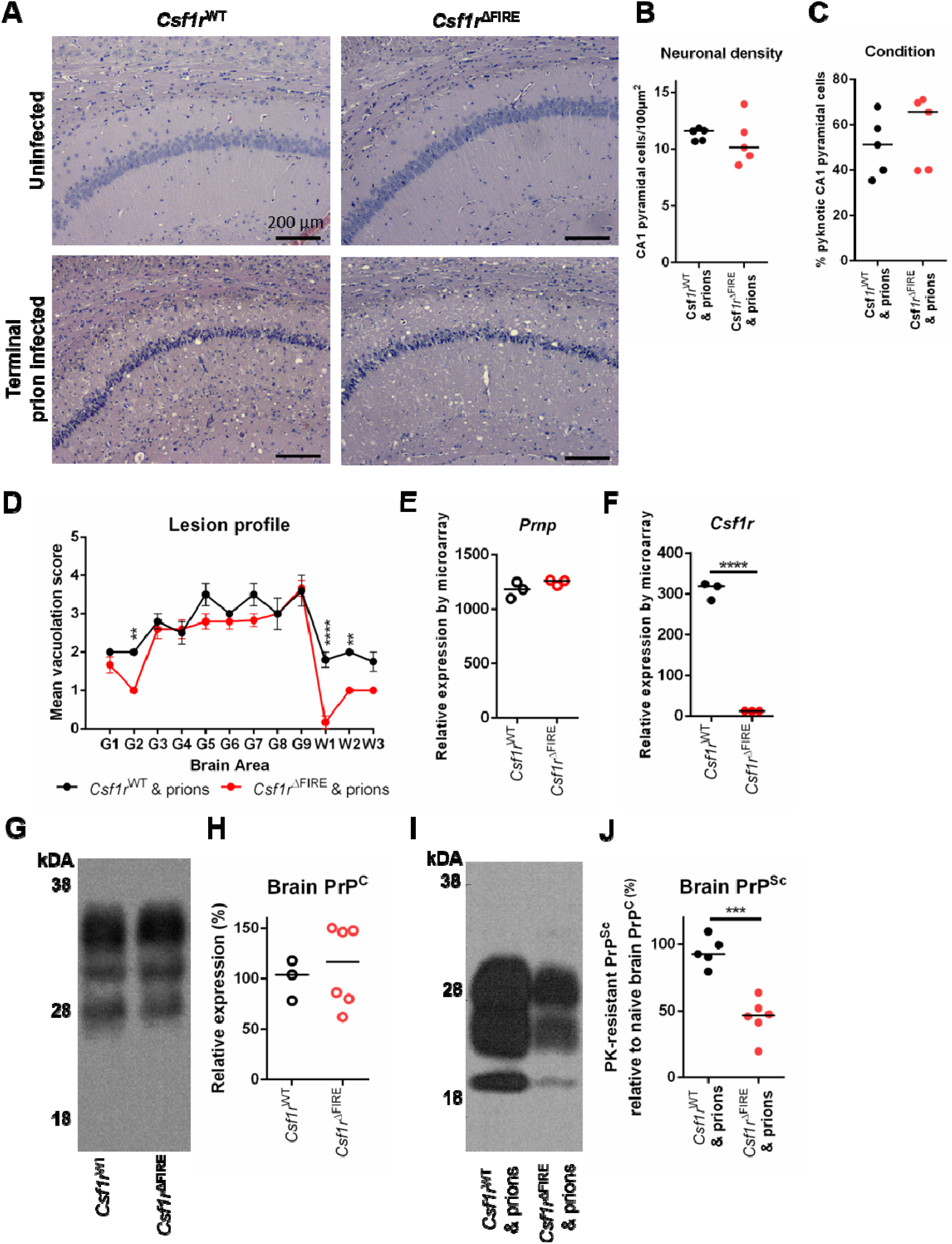
Microglia-deficiency effects on prion-specific vacuolation and prion accumulation. (A) Hematoxylin and eosin stained hippocampus CA1 of uninfected or terminal prion infected *Csf1r*^WT^ or *Csf1r*^ΔFIRE^ mice. Scale bars = 200 µm. (B) Hippocampal CA1 pyramidal cell density in terminal prion infected *Csf1r*^WT^ or *Csf1r*^ΔFIRE^ mice. Points show individual mice, bar = median. Student’s T-test. C) Assessment of neuronal condition expressed as percentage of total neurons pyknotic in terminal prion infected *Csf1r*^WT^ or *Csf1r*^ΔFIRE^ mice. Points show individual mice, bar = median. Student’s T-test. (D) Lesion profile analysis of prion-infected brains N=5-6. Points represent the mean vacuolation score, error bars =D±DSEM. Two-way ANOVA, Sidak’s multiple comparisons test \*\**P* < 0.005, \*\*\*\**P* < 0.0001. (E) Microarray analysis of relative gene expression of *Prnp* in the brain. Points show individual mice, bar = median. Student’s T-test. (F) Microarray analysis of relative gene expression of *Csf1* in the brain. Points show individual mice, bar = median. Student’s T-test \*\*\*\**P* < 0.0001. (G) Western blot analysis of uninfected *Csf1r*^WT^ and *Cs1fr*^ΔFIRE^ mouse brain, probed with anti-PrP antibody clone BH1. Relative protein sizes indicated in kilodaltons (kDa). (H) Quantitation of relative brain PrP^C^ expression in the brains of uninfected *Csf1r*^WT^ and *Cs1fr*^ΔFIRE^ mice. Points show individual mice, bar = median. Student’s T-test. (I) Western blot analysis of terminal prion-infected *Csf1r*^WT^ and *Cs1fr*^ΔFIRE^ mouse brain, probed with anti-PrP antibody clone BH1. Relative protein sizes indicated in kilodaltons (kDa). (J) Quantitation of relative PrP^Sc^ accumulation in the brains of terminal prion-infected *Csf1r*^WT^ and *Cs1fr*^ΔFIRE^ mice. Points show individual mice, bar = median. Students T-test \*\*\**P* < 0.001.

The relative expression level of PrP^C^ can directly influence prion disease duration (Manson et al., 1994, Fischer et al., 1996, Weissmann and Flechsig, 2003). Previous expression profiling of the cortex of *Csf1r*^ΔFIRE^ compared *Csf1r*^WT^ mice revealed no impacts on expression of *Prnp* mRNA (which encodes PrP^C^) or any other neuron-associated transcripts (Rojo et al., 2019). Expression of *Prnp* mRNA in the hippocampus in published mRNA microarray data GEO dataset GSE108207 (Rojo et al., 2019) (**Figure 3E**) was similar despite loss of *Csf1r* expression (**Figure 3F**) and whole brain PrP^C^ protein (**Figure 3G&H**) was similar between naïve *Csf1r*^ΔFIRE^ mice and *Csf1r*^WT^ mice. Partial-deficiency or temporary ablation of microglia during CNS prion infection was reported to accelerate the accumulation of prion-disease-specific PrP^Sc^ in the brain (Zhu et al., 2016, Carroll et al., 2018). By contrast, PrP^Sc^ accumulation was reduced in the brains of *Csf1r*^ΔFIRE^ compared to *Csf1r*^WT^ mice with terminal pathology (**Figure 3I&J**).

### Altered neuropathology in the absence of microglia during CNS prion disease

Consistent with data presented in **Figure 3I&J**, immunostaining for prion disease-associated PrP (PrP^d^) in the brains of *Csf1r*^ΔFIRE^ mice at the terminal stage was approximately 50% of the intensity detected in *Csf1r*^WT^ mice (**Figure 4A&B**). Since the accumulation of PrP^Sc^ within the brain increases as the infection proceeds (Tatzelt et al., 1999), this finding is most likely a consequence of their significantly shortened survival times and implies that microglia deficiency produces hyper-sensitivity to the accumulation of PrP^Sc^.

**Figure 4.**
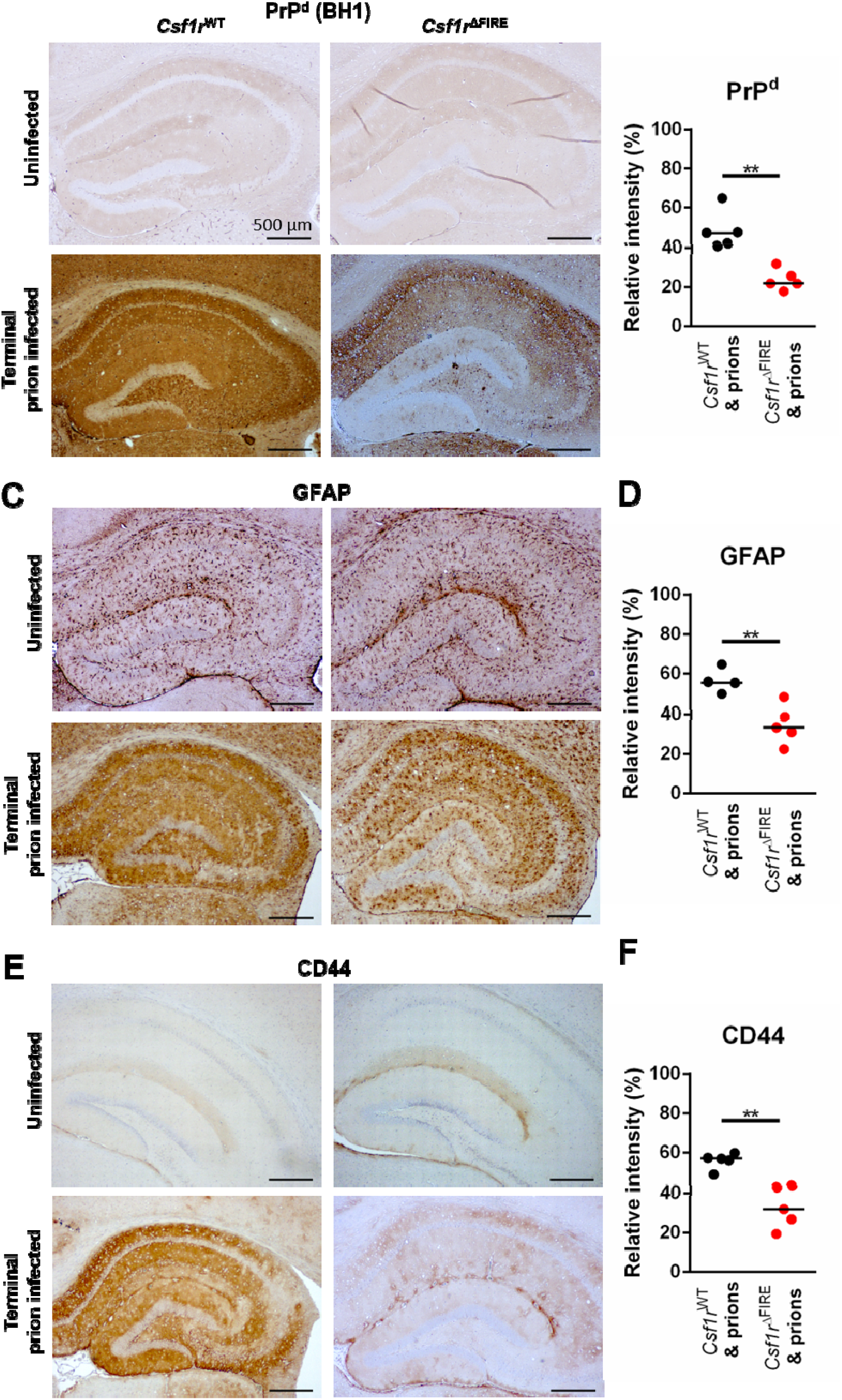
Microglial deficiency reduces terminal neuropathology. (A) Immunohistochemical assessment of PrP^d^ accumulation using anti-PrP antibody clone BH1 in the hippocampus of uninfected or terminal prion infected *Csf1r*^WT^ and *Cs1fr*^ΔFIRE^ mice. DAB (brown) immunostaining lightly counterstained with hematoxylin (blue). Scale bars = 500 µm. (B) PrP^d^ immunostaining quantified by relative intensity. Points show individual mice, bar = median. Student’s T-test \*\**P* < 0.005. (C) Immunohistochemical assessment of GFAP expression in the hippocampus of uninfected or terminal prion infected *Csf1r*^WT^ and *Cs1fr*^ΔFIRE^ mice. DAB (brown) immunostaining lightly counterstained with hematoxylin (blue). Scale bars = 500 µm. (D) GFAP immunostaining quantified by relative intensity. Points show individual mice, bar = median. Student’s T-test \*\**P* < 0.005. (E) Immunohistochemical assessment of CD44 expression in the hippocampus of uninfected or terminal prion infected *Csf1r*^WT^ and *Cs1fr*^ΔFIRE^ mice. DAB (brown) immunostaining lightly counterstained with hematoxylin (blue). Scale bars = 500 µm. (F) CD44 immunostaining quantified by relative intensity. Points show individual mice, bar = median. Student’s T-test \*\**P* < 0.005.

CNS prion disease is accompanied by extensive reactive astrocytosis characterized by high levels of expression of glial fibrillary acidic protein (GFAP), CD44 and the CD44v6 alternative splice variant (Bradford et al., 2019). Microglia and microglial-derived factors have been shown to induce reactive astrocytosis in a range of neurodegenerative conditions (Liddelow et al., 2017, Kunyu Li, 2019, Vainchtein and Molofsky, 2020). Despite the absence of microglia, reactive astrocytes expressing high levels of GFAP (**Figure 4C&D**) and CD44 (**Figure 4E&F**) were increased in the brains of prion-infected *Csf1r*^ΔFIRE^ mice but the level of GFAP^+^ and CD44^+^ immunostaining was lower than in infected *Csf1r*^WT^ mice. As astrocyte activation also increases temporally during CNS prion infection (Hwang et al., 2009, Bradford et al., 2019), this again is most likely a consequence of the *Csf1r*^ΔFIRE^ mice succumbing to terminal prion disease significantly earlier than infected *Csf1r*^WT^ mice. In summary, these data reveal that although CNS prion disease duration is shorter in microglia-deficient *Csf1r*^ΔFIRE^ mice, this is not accompanied by increased neuronal vacuolation, prion accumulation, GFAP or CD44 upregulation at the terminal stage, when compared to infected *Csf1r*^WT^ mice.

### Absence of induction of neurotoxic ‘A1’ or neuroprotective ‘A2’ reactive astrocyte-associated genes in the brains of prion-infected microglia-deficient mice

Reactive astrocytes may be classified into distinct functional subclasses; an A1 subclass with neurotoxic activity and A2 astrocytes considered neurotrophic (Liddelow et al., 2017). Microglia-derived factors have been implicated in the induction of pan- and A1-reactive astrocyte-associated genes (Liddelow et al., 2017). Consistent with the immunohistochemistry data presented in Figure 4, high levels of mRNA encoding the pan-reactive astrocyte-associated genes *Gfap* (**Figure 5A**), *Cd44* (**Figure 5B**) and *Cd44v6* (**Figure 5C**) were detected in the brains of prion-infected *Csf1r*^WT^ mice. LPS-mediated induction of expression of pan-reactive astrocyte-associated genes including *Gfap* and *Cd44* was reported to be blocked in microglia-deficient *Csf1r*^-/-^ mice (Liddelow et al., 2017). However, because of the limited viability of *Csf1r*^-/-^ mice, these studies were performed at postnatal day 8, and these mice are also deficient in peripheral macrophage populations. In the *Csf1r*^ΔFIRE^ mice, the expression of *Gfap, Cd44* and *Cd44v6* was upregulated in response to prion infection despite the complete absence of microglia. These data demonstrate CNS prion-induced reactive astrocyte activation is not dependent on the presence of microglia.

**Figure 5.**
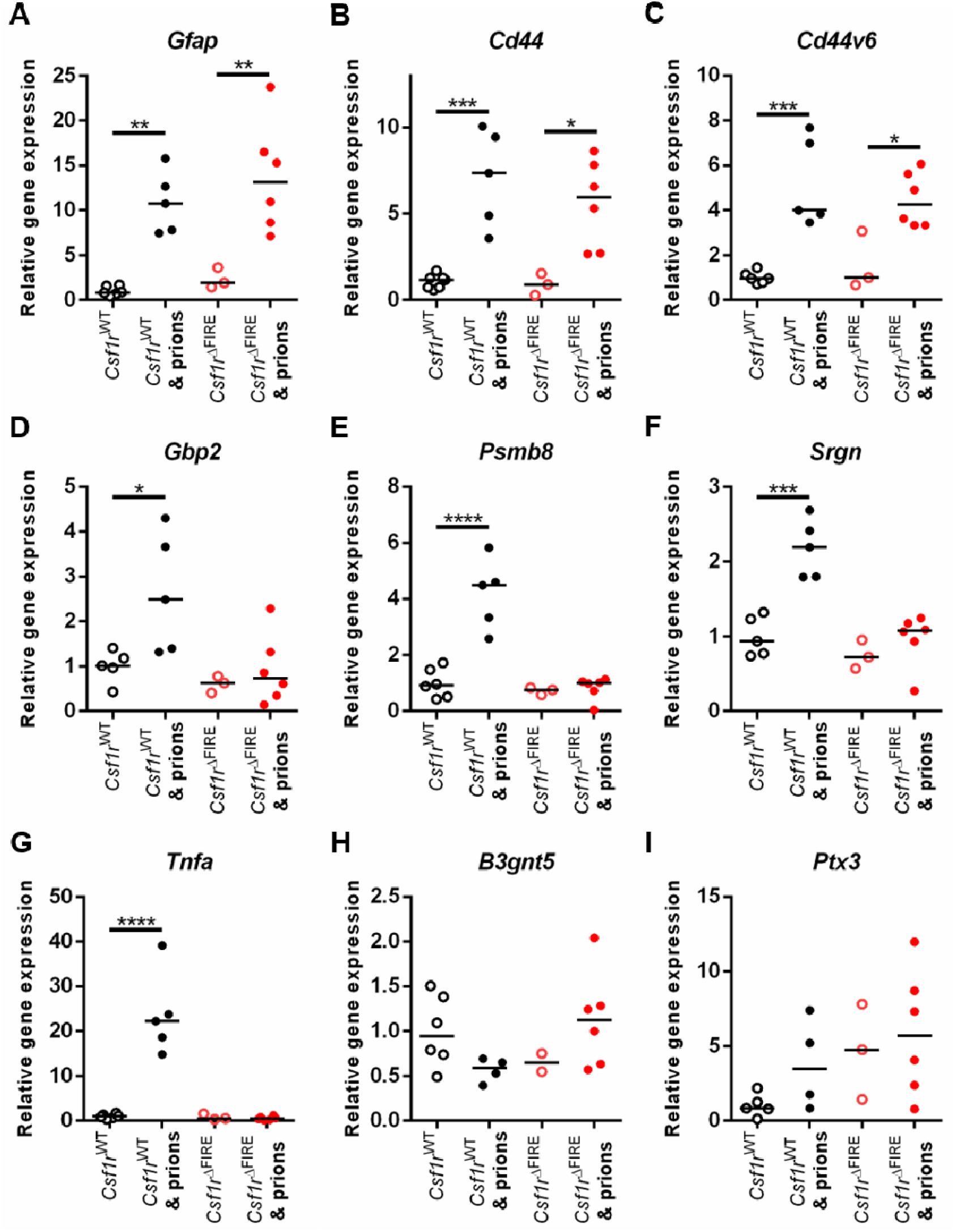
Microglia-deficiency alters astrocyte response to prions. (A) *GFAP* expression analysis via RT-qPCR in uninfected or terminal prion-infected brains from *Csf1r*^WT^ or *Csf1r*^ΔFIRE^ mice. Points show individual mice, bar = median. ANOVA \*\**P* < 0.005. (B) *Cd44* expression analysis. ANOVA \**P* < 0.05, \*\*\**P* < 0.001. (C) *Cd44v6* expression analysis. ANOVA \**P* < 0.05, \*\*\**P* < 0.001. (D) *Gbp2* expression analysis. ANOVA \**P* < 0.05. (E) *Psmb8* expression analysis. ANOVA \*\*\*\**P* < 0.0001. (F) *Srgn* expression analysis. ANOVA \*\*\**P* < 0.001. (G) *Tnfa* expression analysis. ANOVA \*\*\*\**P* < 0.0001. (H) *B3gnt5* expression analysis. ANOVA. (I) *Ptx3* expression analysis. ANOVA.

At the terminal stage of prion disease, the reactive astrocytes display a mixed A1 and A2 transcriptomic signature (Hartmann et al., 2019, Donaldson et al., 2020). The expression of the neurotoxic A1 astrocyte-associated genes *Gbp2, Psmb8* and *Srgn* was upregulated in the brains of terminal prion-infected *Csf1r*^WT^ mice, but absent in *Csf1r*^ΔFIRE^ mice (**Figure 5D-5F**). Microglia-derived cytokines including tumor necrosis factor (TNFα are important inducers of neurotoxic A1 reactive astrocyte activation. Indeed, *Tnf* was elevated in the brains of prion-infected *Csf1r*^WT^ mice but absent in *Csf1r*^ΔFIRE^ mice, coincident with the lack of induction of A1 reactive astrocyte-associated gene expression. Consistent with previous data from the brains of mice infected with ME7 scrapie prions (Donaldson et al., 2020), neuroprotective A2 astrocyte-associated genes (*B3gnt5, Ptx3*) were not induced in the brains of infected *Csf1r*^WT^ or *Csf1r*^ΔFIRE^ mice (Figure 5H, I). Together these data show that CNS prion disease in microglia-deficient *Csf1r*^ΔFIRE^ mice is accompanied by reactive astrocytosis, but lacks evidence of a typical neurotoxic A1 or neuroprotective A2 profile.

### *Csf1r*^ΔFIRE^ mice display accelerated onset of vacuolation but unaltered kinetics of prion accumulation

To determine how disease progression was affected by the absence of microglia, brains were collected from groups of *Csf1r*^WT^ and *Csf1r*^ΔFIRE^ mice at 98 dpi prior to the histopathological detection of neuronal loss. Prion-specific vacuolation was already more severe in prion-infected *Csf1r*^ΔFIRE^ mice in multiple brain regions, including dorsal medulla, superior colliculus, hypothalamus and cerebellar peduncles (**Figure 6A**; vacuolation scoring areas G1, G3, G4, and W3). However, within the hippocampus little evidence of prion-specific vacuolation (**Figure 6A**, vacuolation scoring area G6; **Figure 6C** upper panels) or neuronal loss (**Figure 6B**) was observed in brains from each group at this time. The early ‘synaptic’ patterned PrP^d^ deposition and mild reactive astrocytosis also presented to a similar extent in the hippocampus of infected *Csf1r*^ΔFIRE^ and *Csf1r*^WT^ mice at this time point (**Figure 6C**, middle and lower panels, respectively).

**Figure 6.**
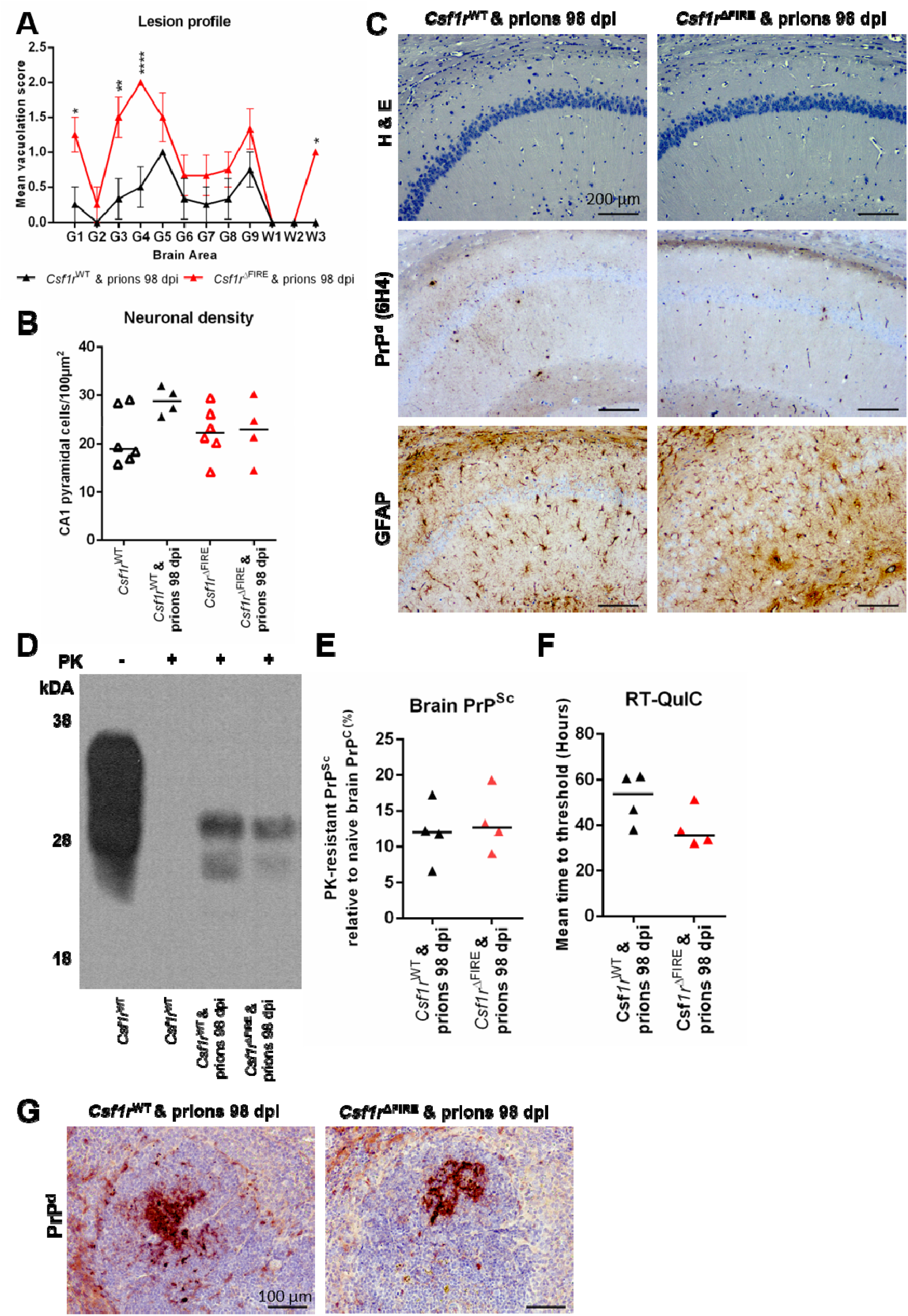
Microglial deficiency accelerates prion vacuolation but not brain or peripheral prion accumulation. (A) Lesion profile analysis of prion-infected brains at 98 dpi. N=4 per group. Points represent the mean vacuolation score, error bars =D±DSEM. Two-way ANOVA, Sidak’s multiple comparisons test \**P* < 0.05, \*\**P* < 0.01, \*\*\*\**P* < 0.0001. (B) Hippocampal CA1 pyramidal neuron density was assessed in uninfected and 98 dpi prion-infected mice. Points show individual mice, bar = median. ANOVA. (C) Hematoxylin and eosin (H &E) stained sections used for vacuolation and neuronal density analyses. Immunohistochemical analysis of PrP^d^ accumulation and GFAP expression in 98 dpi prion-infected *Csf1r*^WT^ and *Csf1r*^ΔFIRE^ hippocampus CA1. Scale bars = 200 µm. (D) Western blot analysis as indicated to determine the relative amount of PrP^Sc^ accumulation in the brains of mice from each groupat 98 dpi with prions. (E) Quantitation of PrP^Sc^ levels in brains of 98 dpi prion-infected *Csf1r*^WT^ and *Csf1r*^ΔFIRE^ mice. Points show individual mice, bar = median. Student’s T-test. (F) Relative prion seeding activities in brains at 98 dpi with prions were quantified *in vitro* by RT-QuIC expressed as mean time to threshold. Points show individual mice, bar = median. Student’s T-test. (G) Immunohistochemical analysis of PrP^d^ accumulation in spleens of prion-infected *Csf1r*^WT^ and *Csf1r*^ΔFIRE^ mice at 98 dpi. PrP^d^ immunostaining (red) counterstained with hematoxylin (blue). Scale bar = 100 µm.

The levels of PrP^Sc^ in the brains of *Csf1r*^ΔFIRE^ or *Csf1r*^WT^ mice at 98 dpi were indistinguishable (**Figure 6D&E**). In parallel, the highly sensitive real-time quaking-induced conversion (RT-QuIC) assay was used to quantify the relative prion seeding activity present within the brains of each group (Atarashi et al., 2011). Consistent with data presented in Figure 6C, the relative levels of prion seeding activity were also similar in the brains of infected *Csf1r*^ΔFIRE^ mice and *Csf1r*^WT^ mice (**Figure 6F**).

After IC injection, some of the infectious prions from the inoculum spread to the spleen via the bloodstream where they accumulate in stromal follicular dendritic cells (FDC) (Brown et al., 1999). Following accumulation within the spleen and other secondary lymphoid organs, the prions can subsequently spread back to the brain (Brown et al., 2012, Brown and Mabbott, 2014). In the absence of peripheral macrophages, the accumulation of prions in secondary lymphoid tissues is enhanced (Beringue et al., 2000, Maignien et al., 2005). Since certain peripheral macrophages will also have been ablated in the previous studies (Zhu et al., 2016, Carroll et al., 2018, Lei et al., 2020) it is plausible that this may have increased the burden of prions in the spleen and other secondary lymphoid organs, and by doing so, enhanced their rate of spread to the brain. However, such an effect was unlikely to responsible for the accelerated prion disease in *Csf1r*^ΔFIRE^ mice, as a similar abundance of prion-specific PrP^d^ was detected on FDC in the spleens of *Csf1r*^ΔFIRE^ mice and *Csf1r*^WT^ mice (**Figure 6G**). This is consistent with the demonstration that spleen macrophage populations are not affected in *Csf1r*^ΔFIRE^ mice (Rojo et al., 2019).

### Accelerated onset of reactive astrocyte activation in the absence of microglia

The increased prion-specific vacuolation in multiple brain regions by 98 dpi (**Figure 6A**), for example within the superior colliculus (vacuolation scoring area G3, **Figure 7A**), was associated with profound astrocytosis in the *Csf1r*^ΔFIRE^ mice. However, at this time no evidence of loss of neuronal nuclear antigen NeuN^+^ neurons was observed within this region in either *Csf1r*^ΔFIRE^ or *Csf1r*^WT^ mice (**Figure 7B**). Instead, increased expression of CD44 (**Figure 7C,F**), and increased frequency of GFAP^+^ morphologically reactive astrocytes (**Figure 7D,G)** was observed in the intermediate grey layer (motor associated area) of the superior colliculus of prion-infected *Csf1r*^ΔFIRE^ mice compared to *Csf1r*^WT^ mice at 98 dpi.

**Figure 7.**
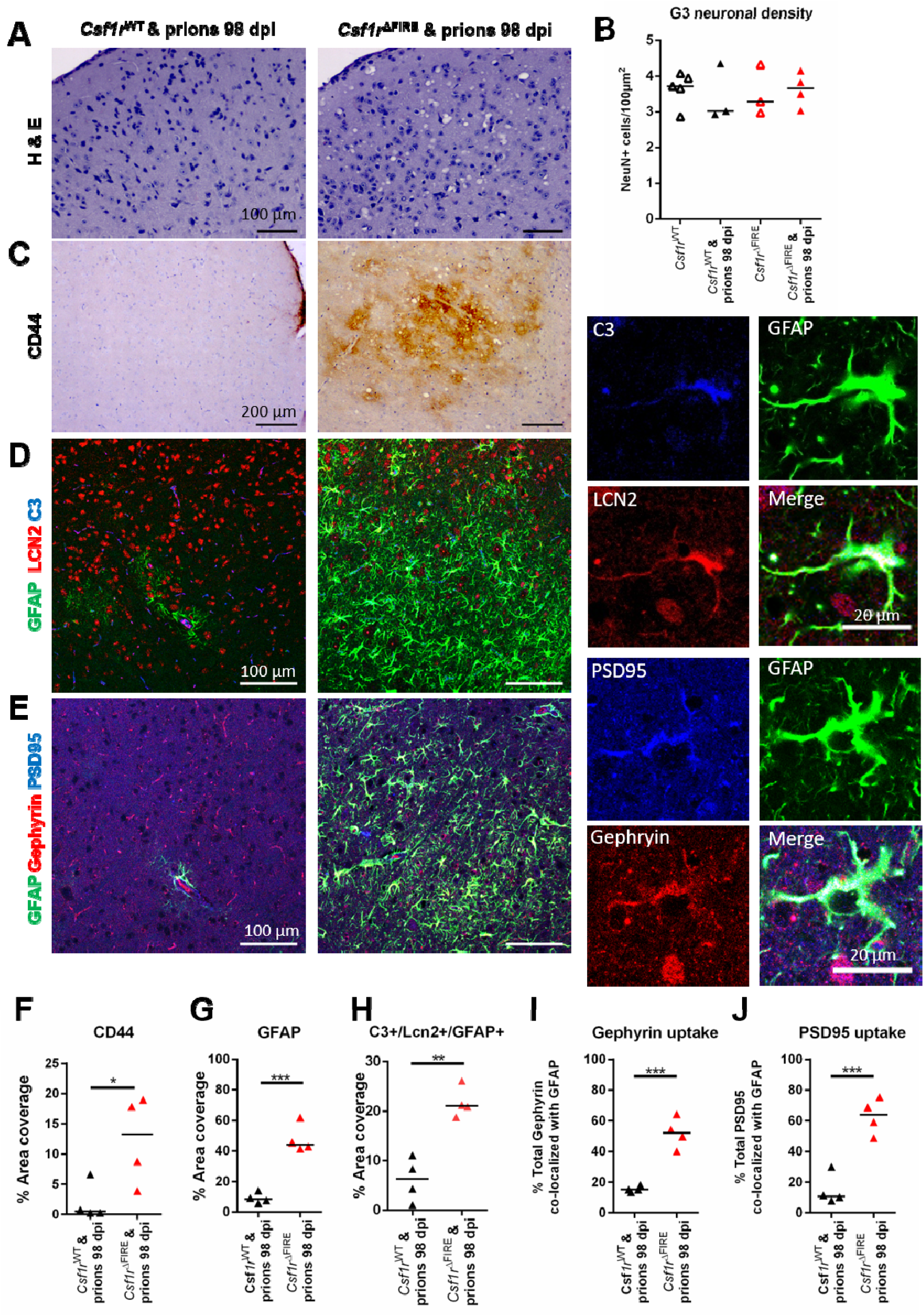
Accelerated astrocyte activation and synaptic pruning in the superior colliculus in the absence of microglia. (A) Hematoxylin and eosin (H & E) stained superior colliculus in 98 dpi prion-infected *Csf1r*^WT^ and *Csf1r*^ΔFIRE^ mice. Scale bars = 100 µm. (B) Superior colliculus neuronal density was assessed via quantitation of the density of NeuN^+^ cells in uninfected and 98 dpi prion-infected *Csf1r*^WT^ and *Csf1r*^ΔFIRE^ mice. Points show individual mice, bar = median. Not significantly different, ANOVA. (C) Immunohistochemical assessment of CD44 expression in 98 dpi prion-infected *Csf1r*^WT^ and *Cs1fr*^ΔFIRE^ superior colliculus. Scale bars = 200 µm. (D) Immunofluorescent assessment of GFAP (green), complement component C3 (blue) and lipocalin2 (LCN2, red) in 98 dpi prion-infected *Csf1r*^WT^ and *Cs1fr*^ΔFIRE^ superior colliculus. Scale bars = 100 µm or 20 µm as indicated. (E) Immunofluorescent assessment of GFAP (green), PSD95 (blue) and gephryin (red) in 98 dpi prion-infected *Csf1r*^WT^ and *Cs1fr*^ΔFIRE^ superior colliculus. Scale bars = 100 µm or 20 µm as indicated. (F) Quantitation of CD44 % area coverage in superior colliculus. Points show individual mice, bar = median. Student’s T-test \**P* < 0.05. (G) Quantitation of GFAP % area coverage in superior colliculus. Points show individual mice, bar = median. Student’s T-test \*\*\**P* < 0.001. (H) Quantitation of C3^+^/LCN2^+^/GFAP^+^ astrocytes. Points show individual mice, bar = median. Student’s T-test \*\**P* < 0.01. (I) Quantitation of gephyrin uptake by astrocytes expressed as % of total gephryin colocalized with GFAP. Points show individual mice, bar = median. Student’s T-test \*\*\**P* < 0.001. (J) Quantitation of PSD95 uptake by astrocytes expressed as % of total PSD95 colocalized with GFAP. Points show individual mice, bar = median. Student’s T-test \*\*\**P* < 0.001.

The innate immune proteins complement component C3 and neutrophil gelatinase-associated lipocalin / lipocalin-2 (NGAL/LCN2) have previously been shown to be upregulated by neurotoxic astrocytes in response to prion infection (Hartmann et al., 2019, Smith et al., 2020, Kushwaha et al., 2021). We observed greater abundance of C3^+^/LCN2^+^/GFAP^+^ morphologically reactive astrocytes within the superior colliculus in *Csf1r*^ΔFIRE^ compared to *Csf1r*^WT^ mice at 98 dpi (**Figure 7D,H**).

Astrocytes in the steady state prune synapses to help maintain neural circuitry (Chung et al., 2013). Abnormal astrocyte synaptic pruning has been implicated in the pathogenesis of some neurodegenerative disorders (reviewed in (Lee and Chung, 2019)), and synaptic alterations are considered to contribute to the early behavioral changes observed in during CNS prion disease (Cunningham et al., 2003). We therefore assessed the localization of the post-synaptic proteins gephyrin and post-synaptic density protein 95 (PSD95) in relation to GFAP^+^ astrocytes (**Figure 7E**). The co-localization of both post-synaptic marker proteins within GFAP^+^ morphologically reactive astrocytes was increased in the superior colliculus of *Csf1r*^ΔFIRE^ compared to *Csf1r*^WT^ mice at 98 dpi (**Figure 7E**). Morphometric analyses suggested over half of the total amount of these synaptic proteins detected in *Csf1r*^ΔFIRE^ mice were within astrocytes (**Figure 7I&J**). These data reveal a statistically significant increase in synaptic engulfment by reactive astrocytes in the brains of prion-infected *Csf1r*^ΔFIRE^ mice compared to *Csf1r*^WT^ mice at 98 dpi within this region.

### Accelerated onset of unfolded protein response in the absence of microglia

Accumulation of misfolded PrP^Sc^ in the brain triggers the unfolded protein response in reactive astrocytes (Smith et al., 2020). Specifically, phosphorylation of protein kinase-like endoplasmic reticulum kinase (PERK) causes the transient shutdown of protein synthesis via phosphorylation of eukaryotic translation initiation factor 2A (eIF2α). Inhibition of PERK-eIF2α signaling in astrocytes alleviated prion-induced neurodegeneration (Smith et al., 2020).

Levels of PERK and eIF2α expression were assessed in brains of uninfected mice and revealed no difference between naive *Csf1r*^WT^ and *Csf1r*^ΔFIRE^ mice (**Figure 8A-C**). In uninfected mice we were unable to detect phosphorylated PERK and eIF2α (**Figure 8A**). However, the levels of phosphorylated PERK and eIF2α were statistically significantly increased in the brains of infected *Csf1r*^ΔFIRE^ mice when compared to infected *Csf1r*^WT^ mice at 98 dpi (**Figure 8D-F**). Immunohistochemical analysis also revealed phosphorylated PERK expression in GFAP^+^ reactive astrocytes and neurons in infected *Csf1r*^ΔFIRE^ mice, particularly within the superior colliculus (**Figure 8G**), coincident with the increased vacuolation and reactive astrocytosis in this region at 98 dpi (**Figure 7**). Conversely, phosphorylated PERK expression in GFAP^+^ reactive astrocytes was not detected in brain regions such as the hippocampus (**Figure 8G**) that displayed little evidence prion-specific vacuolation at this time point (**Figure 6A**).

**Figure 8.**
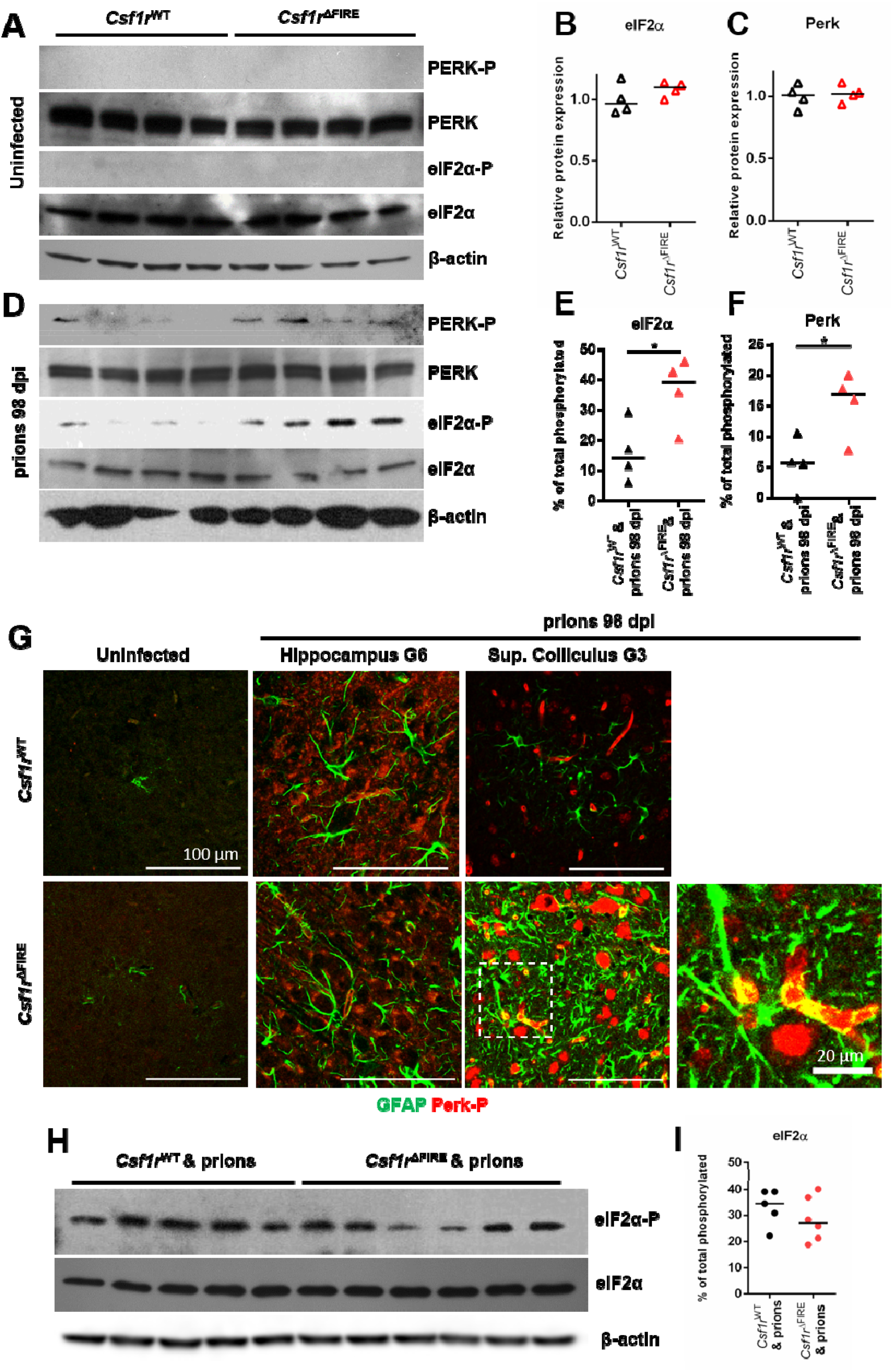
Increased unfolded protein response pathway is associated with earlier astrocyte activation. (A) Western blot analyses of uninfected *Csf1r*^WT^ and *Cs1fr*^ΔFIRE^ mouse brains for unfolded protein response components as indicated, β-actin displayed as a loading control. (B) Quantitation of relative expression levels of eIF2α in uninfected *Csf1r*^WT^ and *Cs1fr*^ΔFIRE^ mouse brains. Points show individual mice, bar = median. Student’s T-test. (C) Quantitation of relative expression levels of PERK uninfected *Csf1r*^WT^ and *Cs1fr*^ΔFIRE^ mouse brain. Points show individual mice, bar = median. Student’s T-test. (D) Western blot analysis of 98 dpi prion-infected *Csf1r*^WT^ and *Cs1fr*^ΔFIRE^ mouse brain for unfolded protein response components as indicated. (E) Quantitation of the percentage of total phosphorylated eIF2α in 98 dpi prion-infected *Csf1r*^WT^ and *Cs1fr*^ΔFIRE^ mouse brain. Points show individual mice, bar = median. Student’s T-test \**P* < 0.05. (F) Quantitation of the percentage of total phosphorylated PERK in 98 dpi prion-infected *Csf1r*^WT^ and *Cs1fr*^ΔFIRE^ mouse brain. Points show individual mice, bar = median. Student’s T-test \**P* < 0.05. (G) Immunohistochemical analysis of phosphorylated PERK (PERK-P;red) and GFAP (green) in uninfected and 98 dpi prion infected *Csf1r*^WT^ and *Cs1fr*^ΔFIRE^ superior colliculus (G3) and hippocampus (G6). Scale bars = 100 µm or 20 µm as indicated. (H) Western blot analysis of terminal prion-infected brain homogenates probed for unfolded protein response components as indicated, β-actin displayed as a loading control. (I) Quantitation of the percentage of total phosphorylated eIF2α in terminal prion-infected *Csf1r*^WT^ and *Cs1fr*^ΔFIRE^ mouse brains. Points show individual mice, bar = median. Student’s T-test.

However, by the terminal stage of prion infection similar levels of phosphorylated eIF2α were detected in the brains of each mouse group despite the *Csf1r*^ΔFIRE^ mice succumbing to clinical prion disease earlier (**Figure 8H&I**). Thus, these data suggest that the earlier astrocyte activation and neuronal vacuolation in the prion-infected *Csf1r*^ΔFIRE^ mice was accompanied by an increased unfolded protein response.

## Discussion

In this study we investigated prion neuropathogenesis in microglia-deficient *Csf1r*^ΔFIRE^ mice. Spongiform vacuolation and neuronal loss at the terminal stage were indistinguishable in *Csf1r*^WT^ and *Csf1r*^ΔFIRE^ mice and the onset of pathology was not correlated with the accumulation of misfolded prions, which are in any case not directly neurotoxic (Benilova et al., 2020). Microglia deficiency did not lead to the increased accumulation of prions in the brain, suggesting that microglial degradation of prions (if it occurs) can be compensated by other cells such as reactive astrocytes. We conclude that the non-redundant function of microglia is to moderate the harmful effects of reactive astrocytes and/or to provide supportive factors to neurons (Sariol et al., 2020). Consistent with that interpretation, microglia can suppress astrocyte phagocytic activity and astrocytes are capable of complete, though slower, clearance of neurons in the absence of microglia (Damisah et al., 2020). Previous studies have used a CSF1R kinase inhibitor to infer the role of microglia in CNS prion disease and reported that overall expression of A1- and A2-reactive astrocyte-associated transcripts in the brain was enhanced upon microglial depletion (Carroll et al., 2018, Carroll et al., 2020). However, use of CSF1R inhibitors can lead to partial depletion of microglia, impact other kinases (e.g. KIT, FLT3), cause localized microglial cell death and impact monocytes and macrophages outside the brain. So, the impacts on pathology should be interpreted with caution (Hume et al., 2020).

During the early stage of prion infection the reactive astrocytes were more abundant in the brains of *Csf1r*^ΔFIRE^ mice. Although there was no induction of A1 neurotoxic astrocyte-associated genes, the reactive astrocytes displayed signs of enhanced pruning of neuronal synapses. The observation of activated astrocytes engulfing synapses in the superior colliculus (G3) region of the brains of *Csf1r*^ΔFIRE^ mice at 98 dpi with prions was coincident with the commencement of overt clinical signs in these mice at this time. These observations strengthen the hypothesis that loss of neuronal connectivity underlies neurological symptoms and precedes complete loss of neurons (Jeffrey et al., 2000, Brown et al., 2001, Cunningham et al., 2003). The engulfment of damaged synapses and neurons by reactive astrocytes could provide a clearance mechanism to protect surrounding undamaged neurons and synapses, as neuronal damage is required for astrocyte-mediated toxicity (Guttenplan et al., 2020).

The reactive astrocytes in the brains of infected *Csf1r*^ΔFIRE^ mice also displayed increased complement C3, LCN2 and phosphorylated activation of PERK and eIF2α in the unfolded protein response pathway. Targeted blockade of this pathway specifically in astrocytes has proved beneficial during prion disease (Smith et al., 2020). Our data from microglia-deficient *Csf1r*^ΔFIRE^ mice indicate that the microglia employ mechanisms to protect the neurons in the brain against prion infection by restricting both phagocytosis and unfolded protein response in astrocytes. A similar role for microglia has recently been described in the suppression of ATP-mediated excitoxicity in neurons (Badimon et al., 2020).

In conclusion, our data indicate that the microglia provide neuroprotection independently of PrP^Sc^ clearance during prion disease and inhibit neurotoxic reactive astrocyte activation. Since astrocytes can contribute to both prion replication (Raeber et al., 1997, Krejciova et al., 2017) and synaptic loss in infected brains, preventing these activities would have therapeutic potential (Smith et al., 2020). Further studies are now required to identify the molecular mechanisms by which the microglia provide neuroprotection during CNS prion disease. The previous characterization of the *Csf1r*^ΔFIRE^ mice included mRNA expression profiling of the hippocampus which identified 85 transcripts that were significantly depleted when compare to wild-type mice, and were presumably not compensated by astrocytes or other cells (Rojo et al., 2019). That list does not include most endosomal and lysosome-associated genes that are more highly expressed by microglia and by inference must be upregulated by other cells in *Csf1r*^ΔFIRE^ mice. An overlapping gene list was generated by expression profiling multiple brain regions in the *Csf1rko* rat (Pridans et al., 2018). Amongst the most down-regulated transcripts are the three subunits of C1q, which have been implicated in regulating astrocyte function (Liddelow et al., 2017, Clarke et al., 2018) and neurodegeneration (Cho, 2019) and have complex roles in neuronal development and homeostasis (Vukojicic et al., 2019). These *Csf1r*-dependent genes provide a short list of non-redundant pathways that may be used by microglia to provide this neuroprotection and restrict the reactive astrocyte activation in prion disease. Paradoxically, given the focus of the literature on harmful functions of microglia, enhancing their functions may provide novel intervention treatments against these devastating neurodegenerative disorders.

## Acknowledgements

We thank University of Edinburgh Biological and Veterinary Services, including Darren Smith & Fraser Laing, Pathology Services Group including Aileen Boyle, The Roslin Institute Histology Suite and Bioimaging facility, including Bob Fleming and Graeme Robertson for helpful advice and technical support. This work was supported by funding from the RS Macdonald Charitable Trust, Edinburgh Neuroscience and project (BB/S005471/1) and Institute Strategic Programme Grant funding from the Biotechnology and Biological Sciences Research Council (grant numbers BBS/E/D/20002173 & BBS/E/D/10002071). For the purpose of open access, the author has applied a Creative Commons Attribution (CC BY) licence to any author accepted manuscript version arising from this submission. The authors declare no scientific or financial conflicts of interest.

